# A single pair of leucokinin neurons are modulated by feeding state and regulate sleep-metabolism interactions

**DOI:** 10.1101/313213

**Authors:** Maria E. Yurgel, Priyanka Kakad, Meet Zandawala, Dick Nassel, Tanja A. Godenschwege, Alex C. Keene

## Abstract

Dysregulation of sleep and feeding has widespread health consequences. Despite extensive epidemiological evidence for interactions between sleep and metabolic function, little is known about the neural or molecular basis underlying the integration of these processes. *Drosophila melanogaster* potently suppress sleep in response to starvation, and powerful genetic tools allow for mechanistic investigation of sleep-metabolism interactions. We have previously identified neurons expressing the neuropeptide leucokinin (Lk) as being required for starvation-mediated changes in sleep. Here, we demonstrate an essential role for Lk neuropeptide in metabolic regulation of sleep. Further, we find that the activity of Lk neurons is modulated by feeding state and circulating nutrients, with reduced activity in response to glucose and increased activity under starvation conditions. Both genetic silencing and laser-mediated microablation localize Lk-mediated sleep regulation to a single pair of Lk neurons within the lateral horn (LHLK) that project near primary sleep and metabolic centers of the brain. A targeted screen identified a critical role for AMP-activated protein kinase (AMPK) in starvation-modulated changes in sleep. Disruption of AMPK function in Lk neurons suppresses sleep and increases LHLK activity in fed flies, phenocopying the starvation state. Taken together, these findings localize feeding state-dependent regulation of sleep to a single pair of neurons within the fruit fly brain and provide a system for investigating the cellular basis of sleep-metabolism interactions.

## Introduction

Dysregulation of sleep and feeding has widespread health consequences and reciprocal interactions between these processes underlie a number of pathologies [1–4]. Sleep loss correlates with increased appetite and insulin insensitivity, while short-sleeping individuals are more likely to develop obesity, metabolic syndrome, type II diabetes, and cardiovascular disease [1,3,4]. Although the neural basis for sleep regulation has been studied in detail, little is known about how feeding state and changes in metabolic function modulate sleep [5,6]. Understanding how sleep and feeding states are integrated may provide novel insights into the co-morbidity of disorders linked to sleep and metabolic regulation.

Animals balance nutritional state and energy expenditure in order to achieve metabolic homeostasis [7,8]. In both flies and mammals, diet potently affects sleep regulation, strengthening the idea that sleep and metabolic state interact [5,6,9]. Starvation leads to sleep loss, or disrupted sleep architecture, presumably to induce foraging behavior, while high-calorie diets have complex effects on sleep depending on macronutrient content [10–13]. Behavioral and physiological responses to changes in feeding state are modulated both by cell autonomous nutrient centers in the brain that sense changes in circulating nutrients and through communication between brain and peripheral tissues [14], yet the neural basis for the integration of sleep and feeding state remain poorly understood.

The fruit fly, *Drosophila melanogaster,* provides a powerful model for investigating sleep regulation. Flies display all the behavioral hallmarks of sleep including extended periods of behavioral quiescence, rebound following deprivation, increased arousal threshold, and species-specific changes in posture [15,16]. Many genetic mechanisms regulating sleep are conserved from flies to mammals. In addition, high throughput systems for sleep analysis in *Drosophila* have led to the identification of both novel and highly conserved sleep genes [17,18]. Further, stimulants including caffeine, amphetamine, and cocaine have been shown to suppress sleep in flies [16,19,20]. Thus, at the molecular, pharmacological, and behavioral levels flies provide a model for studying genetic regulation of mammalian sleep.

A number of genes and neurons that are required for the integration of sleep and feeding states have been identified including core-circadian clock genes, metabolic hormones, and sensory neurons [10, 21–23]. While many identified regulators of sleep-metabolism interactions broadly impact sleep and metabolic function [6], a mutation of the DNA/RNA Binding Protein, *translin* (*trsn*), disrupts starvation-induced sleep suppression without affecting sleep or metabolic regulation under fed conditions. While *trsn* is expressed throughout the nervous system, targeted knockdown in ~30 leucokinin (Lk) neurons phenocopies *trsn* mutants raising the possibility that these neurons are required for the integration of sleep and metabolic state [24].

Here, we identify a single pair of Lk neurons in the lateral horn of the fly brain that are required for the integration of sleep and metabolic state. These neurons project to both sleep and metabolic control centers in the brain and are unique because they do not regulate sleep under fed conditions but are required for starvation-induced sleep suppression. Functional imaging reveals that LHLK neurons have reduced activity in response to glucose application and increased activity under starved conditions. The identification of single neurons that integrate sleep and metabolic state provide a model for investigating the cellular mechanisms regulating the integration of sleep and metabolic state.

## Results

Leucokinin neuropeptide has been implicated in regulation of feeding and circadian activity, but a role in sleep has yet to be identified [25,26]. We measured sleep in fed and starved conditions in flies harboring two different mutations of the *Lk* locus. *Lk^c275^* is a hypomorphic allele containing a Piggy bac element upstream of the *leucokinin* gene transcription start site (Fig 1A) [26]. We also used the CRISPR/Cas9 system to generate a recombinant transgenic line (*Lk^-/-^*^(GAL4)^) with a GAL4 element inserted between base 1 to 7 following the ATG start codon. Lk protein was not detected in the brains of *Lk^-/-^*^(GAL4)^ mutants and there was a reduction in expression in *Lk^c275^* mutants, fortifying the notion that these genetic modifications result in a robust reduction of Lk function (Fig 1B-1D). Sleep on food did not differ between *Lk^c275^* and *Lk^-/-^*^(GAL4)^ lines and *w^11118^* controls, suggesting that Lk is not required for sleep regulation under standard feeding conditions (Fig 1E and 1F). Conversely, control *w^1118^* flies robustly suppressed sleep under starved conditions, while sleep in *Lk^c275^* and *Lk^-/-^*^(GAL4)^ flies did not differ between fed and starved conditions, indicating that Lk is required for starvation-dependent modulation of sleep (Fig 1E and 1F). In agreement with previous findings, control flies display starvation induced hyperactivity (S1A Fig) [21,27]. Starvation did not alter waking activity in *Lk^c275^* and *Lk^-/-^*^(GAL4)^ flies, indicating that Lk is required for both starvation induced changes in sleep regulation and hyperactivity (S1A Fig).

**Fig 1.**
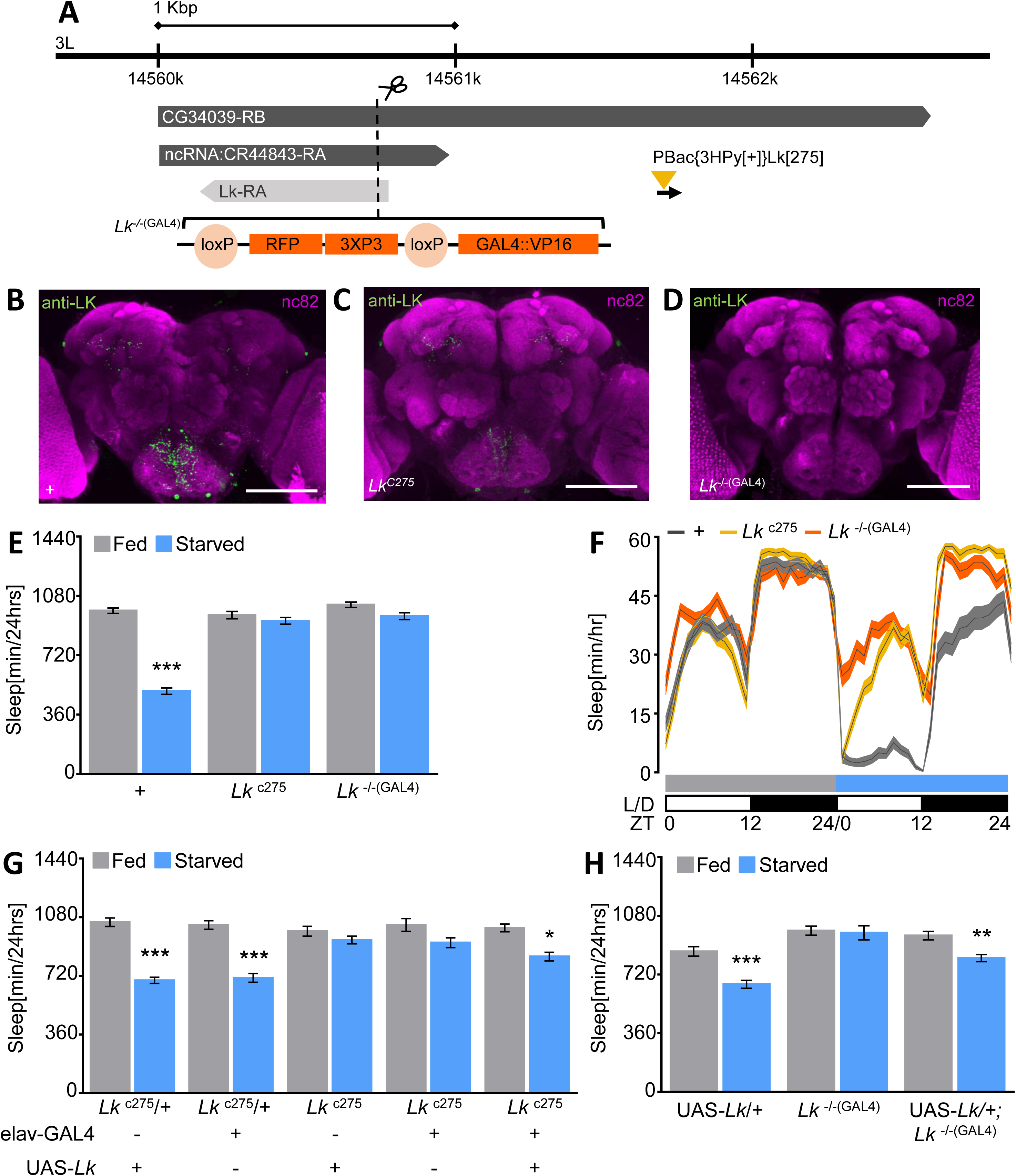
Leucokinin neuropeptide is required for metabolic regulation of sleep. **(A)** The genomic organization of the *Lk* locus. *Lk^c275^* consists of a PiggyBac element inserted 929 base pairs 5’ to the transcription start site of the *leucokinin* gene (gold triangle). The dotted line corresponds to the cleavage site used for Lk^*-/-*(GAL4)^ mutant generation by CRISPR/Cas9-mediated Lk genome engineering. Lk^*-/-*(GAL4)^ contain a GAL4 element replacing base 1 to 7 upstream of the ATG site and a floxed 3xP3-RFP cassette (brackets, orange and light orange). **(B, C, and D)** Immunohistochemistry using Lk antibody in *Lk^c275^* mutants reveals reduction of protein levels **(C)** compared to *w^1118^* control **(B)**, while LK is absent in *Lk^-/-^*^(GAL4)^ muntants **(D)**. The brain was counterstained with the neuropil marker nc82 (magenta). Scale bar = 100μm **(E)** Sleep is significantly reduced in starved *w^1118^* controls (*p*<0.001, n=140), while no significant differences are observed in *Lk^c275^* (n=58, *p*=0.92) or *Lk^-/-^*^(GAL4)^ (n≥47; *p*=0.48) mutant flies. Sleep on food does not differ between control and *Lk^c275^*(*p*=0.87) or *Lk^-/-^*^(GAL4)^ (*p*=0.49), Two-way ANOVA (F(2,486)=58.26). **(F)** Sleep profile of hourly sleep averages over a 48 hour experiment. Flies are placed on food during day 1 (fed, grey), then transferred to agar during day 2 (starved, blue). Sleep does not differ between any of the groups during day 1. *Lk^c275^* and *Lk^-/-^*^(GAL4)^ have increased sleep on agar during day 2. **(G)** Pan-neuronal rescue of *Lk^c275^* (elav-GAL4;*Lk^c275^*>UAS-*Lk*;*Lk^c275^*, n=17, *p*=0.04) restores starvation-induced sleep suppression compared to *Lk^c275^* mutant controls UAS-*Lk/+*;*Lk^c275^*(n=23; *p*>0.99) and elav-GAL4/+;*Lk^c275^* (n=20, *p*=0.70). Sleep duration on agar (starved) does not differ significantly between rescue UAS-*Lk/+*;*Lk^c275^*/+ (n=30, *p*=0.09) or elav-GAL4/*+*;*Lk^c275^*/+ (n=51, *p*=0.11), Two-way ANOVA (F(4,272)=8.97). **(H)** Expression UAS-*Lk* in neurons labeled by *Lk^-/-^*^(GAL4)^ restores starvation induced suppression (n=30, *p*=0.001) compared to *Lk^-/-^*^(GAL4)^ flies (n=15, *p*>0.99). Flies harboring one copy of the UAS-Lk alone (UAS-*Lk*/+) suppress sleep in response to starvation (n=21, *p*<0.0001). There were no significant differences during fed state between control (UAS-*Lk/+*) and rescue flies (*p*=0.14), or *Lk^-/-^*^(GAL4)^ (*p*=0.07), Two-way ANOVA (F(2,126)=5.46). All columns are mean ± SEM; \**p*<0.05; \*\**p*<0.01; \*\*\**p*<0.001.

To verify that the sleep phenotype was caused by loss of *Lk*, we restored *Lk* in the background of each mutant and measured sleep. Pan-neuronal rescue (*Lk^c275^*;elav-GAL4/UAS-*Lk*) restored starvation-induced sleep suppression (Fig 1G) and starvation-induced hyperactivity (S1B Fig), and these flies did not differ from heterozygous controls (*Lk^c275^*;elav-GAL4/+, UAS-*Lk*;*Lk^c275^*/+). Similarly, rescue in Lk neurons (*Lk^-/-^*^(GAL4)^;UAS-*Lk*), where the inserted GAL4 element faithfully drives expression in Lk neurons (S1H Fig), restored starvation-induced sleep suppression (Fig 1H) and starvation-induced hyperactivity (S1C Fig) confirming that Lk is required for the metabolic regulation of sleep. While starvation-induced sleep suppression was abolished in *Lk^c275^* mutants, *Lk^c275^* heterozygous flies suppressed sleep (S1D Fig and S1E Fig). However, flies heterozygous for *Lk^-/-^*^(GAL4)^ failed to suppress sleep in response to starvation (S1D Fig and S1E Fig) and displayed reduced starvation-induced hyperactivity compared to control (S1F Fig), raising the possibility that the phenotype is dominant in CRISPR mutant flies. Expression of a rescue transgene in Lk neurons of flies heterozygous for Lk (*Lk^-/-^*^(GAL4)^/+;UAS-*Lk*), restored starvation-induced sleep suppression (S1G Fig), confirming the specificity of phenotype in *Lk^-/-^*^(GAL4)^ flies.

Leucokinin antibody labels the lateral horn Lk neurons (LHLK) and the subeosophoageal ganglion Lk neurons (SELK), as well as a number of abdominal Lk neurons (ABLK) neurons in the ventral nerve cord (Fig 2A). To localize the population of neurons that regulate starvation-induced sleep suppression, we restricted GAL4 expression primarily to the brain by expressing GAL80, a GAL4 repressor, in ventral nerve-cord using *teashirt*-GAL80 (tsh-GAL80) [28]. Expression of CD8::GFP (Lk-GAL4>CD8:GFP;Lk-GAL80) revealed tsh-GAL80 blocks expression in all but two ventral nerve cord neurons, without affecting expression in the brain (Fig 2B). Silencing the remaining SELK, ALK, and LHLK neurons by expressing light-chain tetanus toxin (TNT) (tsh-GAL80;Lk-GAL4>UAS-TNT) abolished starvation induced sleep suppression, phenocopying the effects of silencing all Lk neurons (Lk-GAL4>UAS-TNT) (Fig 2C and 2D) [29]. Further, no differences in sleep on food were detected between groups, and there was no effect of expressing an inactive variant of tetanus toxin light chain (impTNT) in Lk neurons, fortifying the notion that Lk neurons are dispensable for sleep regulation on food. Taken together, these findings suggest Lk neurons within the brain are required for sleep metabolism interactions.

**Fig 2.**
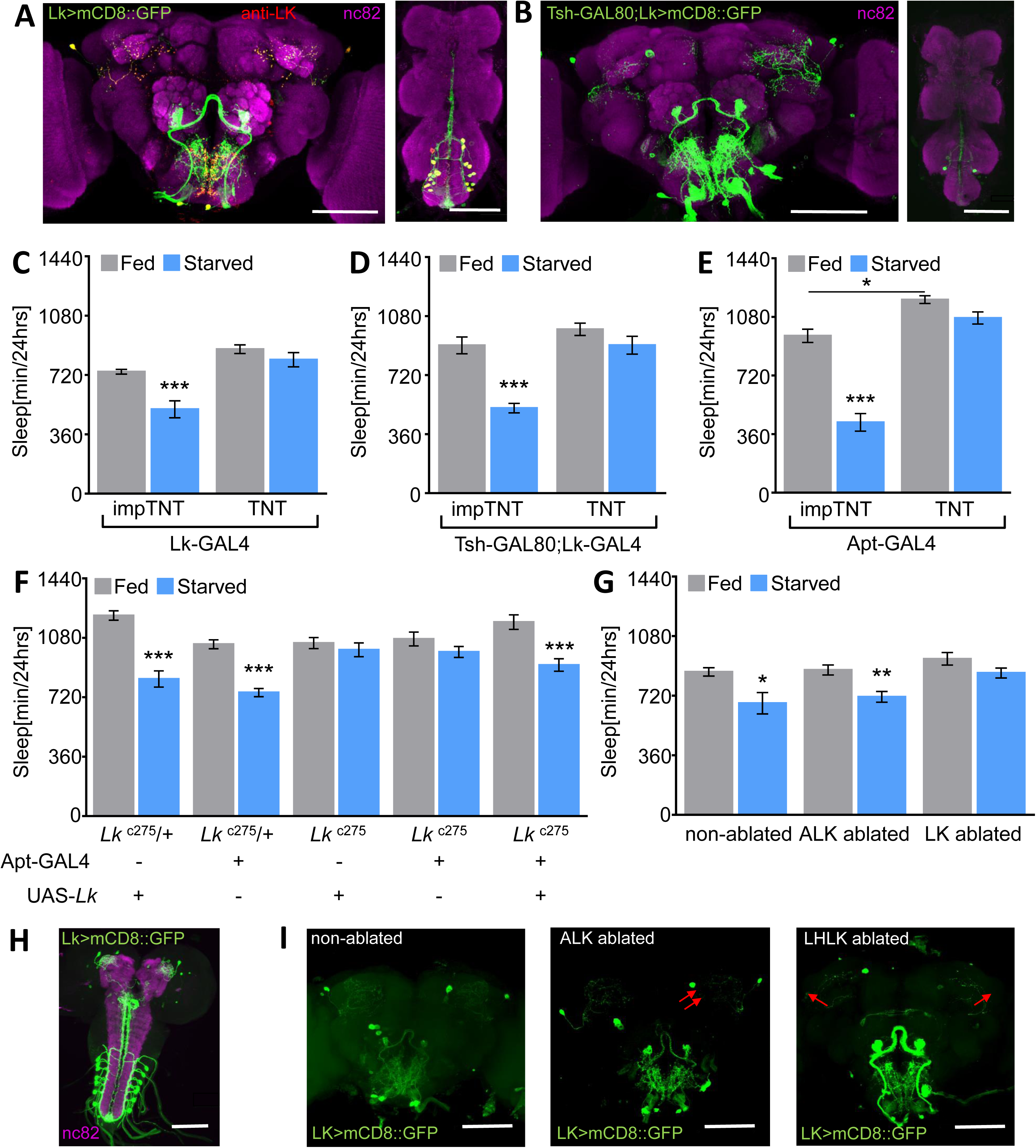
Lateral horn leucokinin neurons are necessary for the metabolic regulation of sleep. **(A)** Whole-brain and ventral nerve cord confocal reconstruction of Lk-GAL4>CD8::GFP. GFP-expressing neurons (green) in the brain labeled one pair of neurons in the subeosphogeal zone, one pair of neurons in the lateral horn, and three pairs of neurons in the medial protorecebrum with variable expression (ALK). In the VNC, Lk is expressed in ~11 pairs of neurons (abdominal Lk neurons; ABLKs). The brain and ventral nerve chord were counterstained with the neuropil marker nc82 (magenta). Scale bar = 100μm **(B)** GFP expression in the ventral nerve chord (ABLK) of flies carrying Lk-GAL4>CD8::GFP is antagonized by tsh-GAL80 transgene. The brain and ventral nerve chord were counterstained with the neuropil marker nc82 (magenta). Scale bar = 100μm **C)** Blocking synaptic release in Lk neurons with TNT impairs starvation induced sleep suppression (n=29, *p*=0.60), while impTNT control supresses sleep (n≥27, *p*=0.0002). No differences were observed between genotypes during fed state (*p*=0.06), Two-way ANOVA (F(1,109)=44.88) **(D)** Starvation induced sleep suppression is absent in tsh-GAL80;Lk-GAL4 flies (n=41, *p*=0.12), while controls expressing inactive impTNT suppress sleep (n=33, *p*<0.001). Sleep duration on food does not differ significantly between tsh-GAL80;Lk-GAL4>TNT and impTNT flies (*p*=0.21), Two-way ANOVA (F(1,144)=22.53). **(E)** TNT expression in Apterous expressing neurons abolishes starvation induced sleep suppression (n≥18, *p*=0.64) and show a significant increase in sleep during fed state (*p*=0.01), compared to impTNT that suppress sleep (n≥34, *p*<0.0001), Two-way ANOVA (F(1,103)=18.22). **(F)** Expression of UAS-*Lk* under control of Apt-GAL4 in the *Lk^c275^* mutant background restores starvation-induced sleep suppression (n=15, *p*=0.002) compared to flies *Lk^c275^* mutant control flies UAS-*Lk*/+;*Lk^c275^* (n=16; *p*>0.99) or Apt-GAL4/+;*Lk^c275^* (n=16, *p*=0.99). Control flies, Apt-GAL4/+;*Lk^c275^*/+ (n≥16, *p*<0.0001) or UAS-*Lk*/+;*Lk^c275^*/+ (n=21, *p*<0.0001), suppress sleep in response to starvation, Two-way ANOVA (F(4, 159)=8.13). **(G)** Flies with bilateral laser ablation of lateral horn leucokinin neurons (LHLK) do not significantly suppress sleep when starved (n=12, *p*=0.09, t=1.18), while ablation of a pair of anterior leucokinin neurons (ALK, n=8, *p*=0.007, t=3.04) and non-ablated controls (n=14, *p*=0.01, t=2.73) suppress sleep in response to starvation, Unpaired t-test. **(H).** Expression pattern of Lk during 3^rd^ instar larval stage visualized with mCD8::GFP. The brain was counterstained with the neuropil marker nc82 (magenta). Scale bar = 100μm. **(I)** Representative images of GFP-expressing Lk neurons post LHLK (right) and ALK ablation (middle), and treated but non-ablated controls (left). Red arrows indicate ablated neurons. Scale bar= 100μm. All columns are mean ± SEM; \**p*<0.05; \*\**p*<0.01; **- \**p*<0.001.

It has previously been reported that *Apterous*-GAL4 drives expression in the LHLK neurons, as well as neurons in the optic lobe, antennal mechanosensory and motor centers (AMMC), and a small population of mushroom body neurons (S2A Fig) [25,30]. Immunostaining with Lk antibody in *Apterous*-GAL4>UAS-CD8::GFP flies revealed co-localization exclusively localized to the LHLK neurons (S2A and S2B Fig). To functionally assess the role of LHLK neurons, we genetically silenced the LHLK neuron, as well as other non-LK cells labeled by *Apterous*-GAL4. Silencing neurons labeled by *Apterous-GAL4* (Apt-GAL4>UAS-TNT) inhibited starvation-induced sleep suppression and promoted sleep on food, while no effects were observed in flies expressing inactive tetanus toxin (Apt-GAL4>UAS-impTNT) (Fig 2E). To verify that the sleep phenotype was due to blocking Lk release from LHLK neurons, we expressed Lk in neurons labeled by Apt-GAL4 in the *Lk^c275^* mutant background and measured sleep. Rescue in Apt-expressing neurons (*Lk^c275^*;Apt-GAL4>*Lk^c275^*;UAS*-Lk*) restored starvation-induced sleep suppression to heterozygote control levels (Apt-GAL4;*Lk^c275^*/+ and UAS-Lk;Lk^c275^/+), whereas flies harboring either GAL4 or UAS in the *Lk^c275^* mutant background failed to suppress sleep (Apt-GAL4;Lk^c275^ and UAS-Lk;Lk^c275^, Fig 2F). These data support a role for the LHLK neurons in starvation-induced sleep suppression, but do not rule out the possibility that other neurons labeled by Apt-GAL4 also contribute to this phenotype.

To verify genetic silencing experiments, we sought to precisely ablate the LHLK neurons and measure their role in starvation-induced sleep suppression. Multi-photon microscopy has been used in diverse genetic models for targeted ablation of neuronal cell types [31–33]. All Lk neurons are present in third instar larvae and labeled by the Lk-GAL4, providing the unique opportunity to independently ablate each subtype and measure the effect on adult behavior in an intact animal (Fig 2H). We selectively induced bilateral ablations of LHLK or unilateral ablation of two control ALK neurons in immobilized 3^rd^ instar with a titanium sapphire multi-photon laser. The ablation of individual neurons could be visualized in larvae as a disruption of GFP-labeled neuronal cell body (S2C Fig). Following ablation larvae were transferred back into food vials and 5-7 day old adult flies were tested for sleep under fed and starved conditions. After behavioral testing, brains were dissected and imaged to verify selective bilateral ablation of the LHLK or unilateral ablation of two ALK neurons (Fig 2I). Flies with ablated ALK neurons suppressed sleep during starvation similarly to controls (Fig 2G). Conversely, bilateral ablation of the LHLK neurons abolished starvation-induced sleep suppression without affecting sleep on food, revealing an essential role for the LHLK neurons in the integration of sleep and metabolic state (Fig 2H-I).

The finding that the LHLK neurons are required for starvation-induced sleep suppression raises the possibility that the activity of Lk neurons is modulated by nutritional state. We selectively expressed a GCaMP6.0-mCherry (UAS-GCaMP-R) fusion protein that allows for ratiometric detection of Ca^2+^ activity [34] in Lk neurons and measured the response to nutrients. The brains of fed controls or 24hr starved flies were imaged for GCaMP6.0 and mCherry signal *ex vivo* (Fig 3A). Flies expressing the GCaMP-mCherry indicator in Lk neurons suppress sleep similarly to control flies harboring Lk-GAL4 alone, indicating that expression of the Ca^2+^ construct does not affect starvation-induced regulation of sleep (S3A Fig). The GCaMP/mCherry ratio was elevated in the LHLK of starved flies compared to fed controls, suggesting these neurons are more active during starvation (Fig 3B). Conversely, no difference in the GCaMP/mCherry ratio between fed and starved state was detected in the SELK neurons that localize to the subesophageal ganglion (Fig 3B). To validate that the activity in Lk neurons is modulated by feeding state in an intact animal, we performed *in vivo* recordings in tethered flies (Fid 3D). Briefly, a portion of the head capsule was removed so that the LHLK neurons were accessible, and the activity was recorded in flies that had been previously fed or starved for 24 hours (Fig 3C). In agreement with *ex vivo* findings, the GCaMP/mCherry ratio was elevated in the LHLK neurons of starved flies, fortifying the notion that Lk neurons are more active during starvation (Fig 3E).

**Fig 3.**
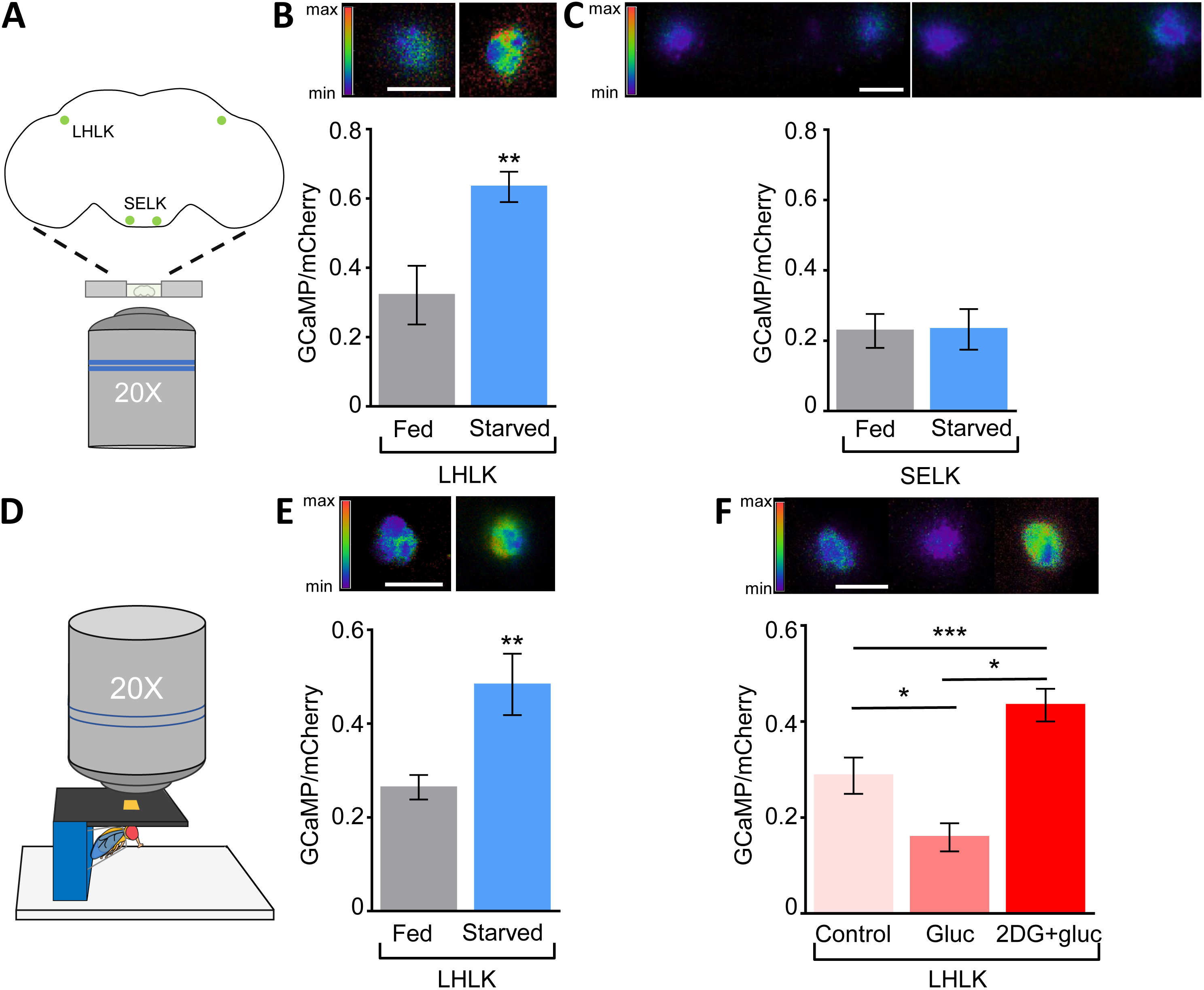
Lateral horn leucokinin neurons have increased activity during starved state. **(A)** Diagram of *ex-vivo* Ca^2+^ imaging. Fed or 24 hr starved adult female flies were dissected and placed dorsally onto chamber. GCaMP/mCherry (UAS-GaMP-R) fluorescence is recorded for 120 seconds with a Ti-Inverted Confocal microscope using a 20X-air objective. **(B and E)** Average ratio of GCaMP6m/mCherry is increased in LHLK neurons during starved state compared to fed state *ex-vivo* (**B**, n≥8, *p*=0.006, t=3.154) and *in-vivo* (**E**, n≥19, *p*=0.006, t=2.89), Unpaired t-test. **(C)** No significant differences in GCaMP6m/mCherry were detected in SELK neurons during fed or starved state *ex-vivo* (n≥7, *p*= 0.95, t=0.06), Unpaired t-test. **(D)** Diagram of *in-vivo* Ca^2+^ imaging. Cuticle of fed or 24 hr starved adult female fly is removed and GCaMP/mCherry fluorescence is recorded for 120 seconds. **(F)** *Ex-vivo* application of 400mM 2DG and 200mM glucose to fed fly brains (n=12) increases the GCaMP6m/mCherry fluorescence ratio in LHLK neurons compared to control (hemolymph-like solution alone, n=11, *p*=0.01) and glucose application (200mM, n=12, *p*<0.0001). Glucose application alone reduced the GCaMP6m/mCherry ratio compared to haemlymph-like solution control (*p*=0.03). One-way ANOVA, (F(2,32)=17.10). Fluorescence intensity scale represents the ratio range of GCaMP6m/mCherry ranging from 2 (max) to 0 (min). Scale bar = 10μm. Error bars for Gcamp6m/mCherry ratio during fed vs starved state indicate SEM; \**p*<0.05; \*\**p*<0.01; \*\*\**p*<0.001.

In mammals, the activity of sleep and wake promoting neurons are directly modulated by glucose and other circulating nutrients [35,36]. It is possible that the activity of Lk neurons is modulated in accordance with feeding state by sensory detection of tastants, or indirectly result from detection of changes in circulating nutrients. To differentiate between these possibilities, the brains of fed flies were removed and treated with either glucose, or the competitive inhibitor of glycolysis, 2 deoxy-glucose (2DG) [37–39]. Application of glucose reduced Ca^2+^ activity in Lk neurons compared to controls treated with *Drosophila* hemolymph-like solution alone, suggesting these neurons are sensitive to circulating glucose (Fig 4E). Further, the combined application of 2-DG and glucose increased Ca^2+^ activity to levels greater then hemolymph alone, mimicking the starved state. Taken together, these findings indicate that the activity of Lk neurons are modulated in accordance with nutrient availability and support the notion that the LHLK neurons are more active during starvation, thereby suppressing sleep.

**Fig 4.**
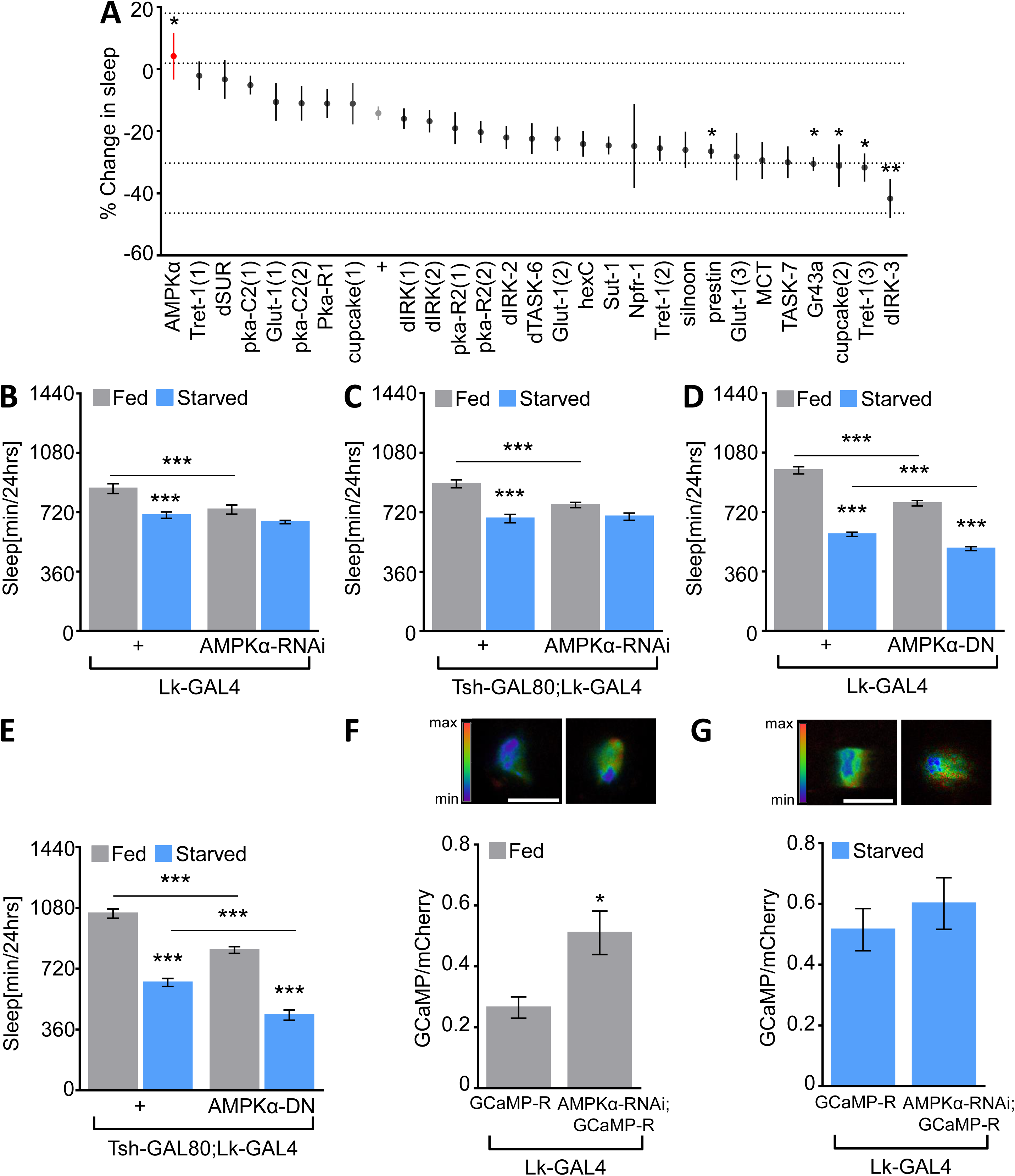
AMPkα is a key nutrient sensor that functions in LHLK. **(A)** The percentage change in sleep in an RNAi screen targeting nutrient sensors or signalling pathway molecules in Lk-GAL4 neurons. Greater sleep suppression was observed in controls Lk-GAL4 and the isogenic host strain for the Vienna *Drosophila* Resource Center RNAi library (Lk-GAL4/+, n=56) compared to AMPKα-RNAi (n=9, *p*=0.04), One-way ANOVA with Dunnett, F(28, 358)=4.47). Dashed lines indicate control mean ±2 SD. **(B)** Knock down of AMPKα in Lk neurons (Lk-GAL4, n=32, *p*=0.09) abolishes starvation induced sleep suppression, while control flies Lk-GAL4/+ suppress sleep (n=30, *p*<0.0001). During fed state, sleep is significantly reduced in Lk-GAL4>UAS-AMPKα-RNAi compared to control (*p*=0.0009), Two-way ANOVA (F_treatment_(1,120)=26.50; F_genotype_(1,120)=13.48). **(C)** Flies expressing UAS-AMPKα-RNAi in brain Lk neurons (tsh-GAL80;Lk-GAL4) fail to suppress sleep in response to starvation (n≥70, *p*=0.06) compared to control flies that suppress sleep (tsh-GAL80;Lk-GAL4/+, n≥50, p<0.0001). Sleep is significantly reduced in tsh-GAL80;Lk-GAL4 >UAS-AMPKα-RNAi compared to control (*p*=0.001), Two-way ANOVA, (F(1,247)=10.66). **(D and E)** Flies expressing a dominant negative form of AMPKα in Lk neurons (**D**, Lk-GAL4, n=89, *p*<0.0001) and brain Lk neurons (**E**, tsh-GAL80;Lk-GAL4, n≥52, *p*<0.0001) significantly suppress sleep in response to starvation. During fed state sleep is significantly reduced in Lk-GAL4> UAS-AMPKα-DN and tsh-GAL80;Lk-GAL4>UAS-AMPKα-DN compared to controls expressing a wild-type copy of AMPKα in all Lk neurons (n=73, *p*<0.0001) or brain Lk neurons (n≥24, *p*<0.0001), Two-way ANOVA, (**D**, F (1,320)=12.25) and (**E**, F(1,168)=0.17). **(F)** *In-vivo* Ca^2+^ imaging during fed state increases the average ratio of GCaMP6m/mCherry in LHLK neurons expressing UAS-AMPKα-RNAi;UAS-GCaMP-R (n=10) compared to flies harboring GCaMP-R alone (n=8, *p*=0.01, t=2.86), Unpaired t-test. **(G)** During starved state no significant differences in LHLK Ca^2+^ activity are observed in Lk-GAL4>UAS-AMPKα-RNAi;UAS-GCaMP flies (n=8) compared to flies harboring GCaMP-R alone (n=8, *p*=0.44, t=0.78), Unpaired t-test. Fluorescence intensity scale represents the ratio range of GCaMP6m/mCherry ranging from 2 (max) to 0 (min). Scale bar = 10μm. Error bars for Gcamp6m/mCherry ratio during fed vs starved state indicate SEM; \**p*<0.05; \*\**p*<0.01; \*\*\**p*<0.001.

The localization of nutrient-dependent changes in activity to LHLK neurons raises the possibility that cell-autonomous nutrient sensors or signaling pathways function within Lk neurons to modulate sleep. To identify regulators of sleep that modulate activity of Lk neurons, we expressed RNAi targeted to 28 RNAi lines encoding putative nutrient sensors or signaling pathways using Lk-GAL4 and measured starvation-induced changes in sleep (Fig 4A). RNAi knockdown of AMPKα Activated Protein Kinase in Lk neurons alone abolished starvation-induced sleep suppression compared to GAL4 controls crossed to the isogenic host strain for the RNAi library (Fig 4A and 4B) [40]. In addition, genetically restricting AMPK knockdown in flies harboring tsh-GAL80 (tsh-GAL80;Lk-GAL4>AMPKα-RNAi) also impaired starvation-induced sleep suppression (Fig 4C). Of particular interest is the finding that knockdown of AMPKα in Lk neurons (Lk-GAL4> AMPKα-RNAi) reduces sleep in fed flies rather than blocking the starvation induced sleep suppression, supporting the notion that inhibition of AMPK in Lk neurons alone is sufficient to induce changes in sleep that are indicative of the starved state. To validate findings obtained with RNAi, we expressed a dominant negative variant of AMPK (AMPKα-DN) in Lk neurons. Similar to the phenotypes obtained with RNAi, flies expressing AMPKα-DN in all Lk neurons (Lk>AMPKα-DN) or primarily brain-restricted Lk neurons (tsh-GAL80;Lk-GAL4>AMPKα-DN) slept less than control flies expressing wild-type AMPK (AMPKα) (Fig 4D,E), indicating that AMPK functions in Lk neurons to promote sleep.

To determine whether inhibition of AMPK signaling changes the physiology of Lk neurons to resemble a starved-like state, we genetically expressed AMPKα-RNAi under control of Lk-GAL4 and measured neuronal activity using UAS-GCaMP-R (Lk-GAL4>UAS-AMPKα-RNAi; UAS-GCaMP-R). Genetic disruption of AMPKα increased Ca^2+^ activity in LHLK neurons of fed flies compared to flies expressing UAS-GCaMP-R alone (Fig 4F). The increase phenocopies changes found in starved control flies, suggesting loss of AMPK increases the activity of Lk neurons, thereby suppressing sleep (Fig 4G). Together these findings suggest AMPK is active within LHLK neurons during the fed state, and reduced AMPK signaling during starvation increases LHLK activity, thereby suppressing sleep.

## Discussion

Our findings reveal that Lk neurons and Lk neuropeptide are selectively required for the integration of sleep and metabolic state. Silencing of Lk neurons, or mutation of the Lk locus does not affect sleep under fed conditions but abolishes starvation-induced sleep suppression. Previous studies have identified a number of genes required for starvation-induced changes in sleep or locomotor activity, yet many of these genes have pleiotropic functions on behavior or metabolic function [21,27,41,42]. For example, the glucagon-like adipokinetic hormone (AKH) is responsible for energy mobilization, and genetic disruption of AKH induces obesity and abolishes starvation-induced hyperactivity [27,43,44]. Similarly, the circadian transcription factors *clock* and *cycle* are required for starvation-dependent regulation of behavior and loss of function affects sleep both in fed and starved conditions [21]. Conversely, neuropeptide F functions within a subpopulation of circadian neurons and is selectively required for metabolic regulation of sleep [23]. Similarly, we find that *Lk* mutants and silencing of Lk neurons has little effect on sleep under fed conditions but disrupts starvation-induced modulation of sleep. These findings suggest that different neural mechanisms regulate sleep under basal conditions and in response to environmental perturbation.

The failure of *Lk* mutants to suppress sleep under starved conditions phenocopies mutation of the RNA/DNA binding protein *trsn.* Loss of *trsn* does not impact feeding behavior but impairs starvation-induced sleep suppression suggesting that *trsn* is not generally required for hunger-induced behavior [45]. While *trsn* is broadly expressed in the fly nervous system [46,47], we previously found that selective knockdown of *trsn* in Lk neurons disrupted starvation induced sleep suppression [45]. These findings raise the possibility that *trsn* functions to regulate changes in physiology of Lk neurons to modulate sleep under starved conditions. In mice, *trsn* is required for dendritic trafficking of brain-derived neurotrophic factor (*BDNF*) mRNA and hippocampus-dependent memory formation suggesting a critical role in synaptic function and plasticity [48,49]. Both sleep loss and starvation affect synaptic architecture and physiology [50–54], raising the possibility that *trsn* is required for state-dependent modulation of synaptic function. *Translin* also functions as a key component in the RNAi silencing complex (RISC) [55–57]. In this context, *trsn* might post-transcriptionally regulate Lk translation, cellular localization, or secretion. Therefore, it is possible that *trsn* may directly modulate Lk function or may be indirectly required for the function of Lk neurons.

Leucokinin is expressed in 4 pairs of bona fide neurons in the brain and 11 pairs in the ventral nerve cord, that regulate diverse behaviors and physiological processes [58–60]. The brain and ventral nerve cord Lk neurons all have distinct projection patterns suggesting unique functions [59]. The ABLK neurons in the ventral nerve cord, have axon terminations in the lateral heart nerves and abdominal ganglion [58] and have been implicated in response to various stressors including starvation, desiccation, and ionic stress [60]. The SELK neurons, in the brain, connect the gustatory receptors to the subesophageal ganglia and ventral nerve cord. Although a specific function has not been identified to SELK neurons, silencing of all Lk neurons disrupts gustatory behavior and a mutation in the *Lk* locus affects meal size [26,61], raising the possibility that these behaviors are regulated by SELK neurons. Lastly, the LHLK neurons project to the superior lateral protocerebrum, medial protocerebrum, and peduncle and axonal stalk of the mushroom bodies [59]. Disruption of Lk within these neurons attenuates circadian rhythms and this effect is localized to the same LHLK neurons that modulate starvation-induced sleep suppression [25]. It is possible that the clock inputs that confer rhythmicity to Lk neurons are overridden by feeding-fasting patterns.

The *Drosophila* genome encodes for a single Lk target, the leucokinin receptor (LkR), that is expressed in the lateral horn, the ventral nerve cord, and the sleep-promoting fan-shaped body [25,26,62]. The fan-shaped body, a suregion of the *Drosophila* central complex, is a primary sleep-promoting region [63,64] raising the possibility that Lk neurons signal to the fan-shaped body to promote sleep or wakefulness during starvation. Supporting this notion, functional analysis suggests Lk-LkR connectivity is proposed to be inhibitory [25,65]. Therefore, the increased activity within LHLK neurons that we report during starvation may inhibit sleep-promoting fan-shaped neurons, resulting in starvation-dependent sleep suppression. Functional imaging of the fan-shaped body in flies with ablated LHLK neurons will provide information about the downstream circuitry through which LHLK neurons suppress sleep under starved conditions.

In *Drosophila,* a number of circulating nutrients including fructose, trehalose, and glucose have been found to affect central brain physiology and behavior [39,66,67]. While nutrients may be detected by gustatory receptors expressed in the periphery to regulate sleep [10,68], sugar receptors and transporters are also expressed in the brain [69]. The identification of LHLK neurons as being active under starvation conditions and suppressed by glucose provide a system to investigate feeding-state dependent changes in neural activity. A number of neurons in the fly brain are acutely regulated by feeding state including the starvation-active Taotie neurons that inhibit insulin producing cells (IPCs) of the pars intercerebralis to regulate insulin-like peptide release under nutrient deprivation conditions [70–72]. Conversely the IPCs, which are cell autonomous nutrient sensors themselves, are activated by glucose through the inhibition of K_ATP_ channels, supporting the notion that ingested nutrients are directly sensed by neurons [73]. Further, the LHLK nutrient phenotype is similar to the neurons within the ellipsoid body labeled by the sodium/glucose co-transporter SLC5A11 that are active during starvation and promote feeding [38,74]. SLC5A11 and its cognate neurons are required for a variety of hunger-induced feeding behaviors, but the effect on sleep has not been identified [74]. Our screen found that knockdown of SLC5A11 in Lk neurons did not affect starvation-induced sleep suppression suggesting alternative regulators of sleep. The identification of LHLK neurons as starvation-active neurons provides a system for identification of additional nutrient sensors that regulate sleep.

The finding that the activity of LHLK neurons are modulated by feeding state have parallels with sleep-regulating neurons in mammals. In mice, both sleep and wake-promoting neurons sense changes in nutrient availability, through direct detection of circulating glucose or hormonal cues [75,76]. For example, glucose inhibits K_ATP_ channels within the sleep-promoting VLPO neurons of the hypothalamus, promoting slow wave sleep [35]. Conversely, hypothalamic neurons expressing the wake-promoting neuropeptide orexin/hypocretin (HCRT) are inhibited by glucose and leptin while activated by the hunger hormone, ghrelin [77,36,78]. Therefore, the findings that LHLK neurons have increased activity during starvation which results in wakefulness suggest that Lk neurons may play a role analogous to orexin/HCRT neurons in mammals.

AMP-activated kinase functions as a cell-autonomous regulator of energy allocation and induces physiological changes associated with starvation [79,80]. AMPK consists of a heterotrimeric complex that is activated by AMP and modulates diverse intercellular signaling pathways including mTOR, FoxO, and SIRT1[81]. Canonically, AMPK is activated during starvation and increases neuronal activity, though this effect varies by neuronal subtype [82,83] For example, in *C. elegans*, starvation-induced AMPK activation leads to inhibition of neurons that modulate local search behavior in response to food deprivation, while promoting activity in neurons that trigger dispersal behavior [84]. Here, we use a dominant negative variant of AMPK with a mutation in the catalytic domain of the alpha subunit to selectively disrupt AMPK function in Lk neurons [82]. Ubiquitous disruption of AMPK in *Drosophila* induces hypersensitivity of the locomotor response to starvation and reduces starvation resistance [82]. Conversely, we find that selectively disrupting AMPK function in Lk neurons promotes starvation-induced hyperactivity and sleep loss during fed state, revealing neural circuit-specific function for AMPK. Further, Ca^2+^ imaging reveals that knockdown of AMPK increases neural activity, mimicking the starvation state. While our findings indicate that AMPK directly modulates the activity of Lk neurons, it is also possible that AMPK modulates Lk transcription or release. The findings that the activity of SELK neurons is not elevated during starvation, raises the possibility of neuron-specific AMPK function. The identification of AMPK as a critical modulator of LHLK neuronal activity and state dependent changes of activity within LHLK neurons provides a system for identifying novel nutrient-sensing and signaling mechanisms that modulate sleep. Further investigation of feeding state dependent changes in Lk signaling, and the identification of neuronal inputs and targets of LHLK neurons will provide mechanistic insight into how animals integrate sleep and metabolic state.

## Methods

### *Drosophila* maintenance and Fly stocks

Flies were grown and maintained on standard food (Bloomington Recipe, Genesee Scientific). Flies were kept in incubators (Powers Scientific; Dros52) at 25°C on a 12:12 LD cycle with humidity set to 55-65%. The background control line used in this study is *w^1118^* fly strain, and all experimental flies were outcrossed 6-8 generation into this background. The following fly strains were ordered from Bloomington Stock Center, w^1118^(5905; [85]), Lk^c275^(16324;[26]), elav-GAL4(8765; [86]), Apterous-GAL4(3041;[87]), UAS-TNT(28996; [29]), UAS-impTNT (28840; [29]), UAS-mCD8::GFP (32186; [88]), UAS-AMPKα (32108; [89]), UAS-AMPKα-DN (32112; [82]). The following lines were generated in this study, Lk^*-/-*(GAL4)^ and UAS-*Lk*. UAS-GCaMP-R and Lk-GAL4 were a kind gift from Greg Macleod and Young-Joon Kim, respectively. tsh-GAL80 [28] was provided by Julie Simpson. *Drosophila* lines used in the RNAi screen originate from the VDRC library [40] and are described in (Table 1).

**Table 1:** Fly strains used in the Screen.

| **Name** | **CG** | **Vienna Stock center ID** | **FlyBase ID** |
| --- | --- | --- | --- |
| GD control |  | V6000 |  |
| cupcake | CG8451 | V48987(1)<br>v3424 | FBgn0031998 |
| npfr1 | CG1147 | v9605 | FBgn0037408 |
| Gr43a | CG1712 | v39518 | FBgn0041243 |
| Glut 1 | CG43946 | v47179(2)<br>v47178(3)<br>v13326(1) | FBgn0264574 |
| Sut-1 | CG5772 | v9950 | FBgn0028563 |
| Tret-1 | CG30035 | v52360(3)<br>v52361(2)<br>v8126(1) | FBgn0050035 |
| TASK-6 | CG9637 | v9073 | FBgn0038165 |
| TASK-7 | CG9361 | v8565 | FBgn0037690 |
| dIRK | CG44159 | v28430(1)<br>v28431(2) | FBgn0039060 |
| dIRK-2 | CG4370 | v4341 | FBgn0039081 |
| dIRK-3 | CG10369 | v3886 | FBgn0032706 |
| dSUR | CG5772 | v6750 | FBgn0025710 |
| Hex-C | CG8094 | v35337 | FBgn0001187 |
| AMPKα | CG3051 | v1827 | FBgn0023169 |
| PKA C2 | CG12066 | v30658(1)<br>v30685(2) | FBgn0000274 |
| PKA R1 | CG42341 | v26328 | FBgn0000275 |
| PKA R2 | CG15862 | v39437(1)<br>v39436(2) | FBgn0000353 |
| Prestin | CG5485 | v5341 | FBgn0036770 |
| MCT | CG8028 | v9163 | FBgn0031010 |
| Silnoon | CG8271 | v4607 | FBgn0033657 |

### Generation of GAL4 knock-in mutants and UAS-Lk

*Lk^-/-^*^(GAL4)^ was generated by Wellgenetics (Taipei City, Taiwan) using the CRISPR/Cas9 system to induced homology-dependent repair (HDR) using one guide RNA (GATCTTTGCCATCTTCTCCAG). At gRNA target site a donor plasmid was inserted containing a GAL4::VP16 and floxed 3xP3-RFP cassette [90]. Following the ATG start site bases 1 to 7 were replaced by the knock in cassette. All lines were generated in the *w^1118^* background [85]. The insertion locus for both mutations was validated by genomic PCR.

### UAS-*Lk*

The full-length open reading frame of Leucokinin was amplified from the Leucokinin-pOT2 plasmid (Drosophila Genomics Resource Center, #1378621) using specific primers (Forward primer: GCCTTTGGCCGTCAAGTCTA and Reverse primer; CTCCAAGTACCGCAGGTTCA) generated by Integrated DNA Technologies, IDT. Amplified sequence was inserted into the pENTER vector (Invitrogen) via TOPO cloning and subsequently recombined into pTW destination vector (DGRC, #1129) using standard gateway cloning protocol as per manufacturer’s instructions (Invitrogen). The plasmids were verified by sequencing (Genewiz LLC). Transgenic lines were established via phiC31**-**mediated integration at the attp40 landing site [91] on the second chromosome (BestGene Inc).

### Behavioral analysis

The *Drosophila* Activity Monitor System (DAMS) detects activity by monitoring infrared beam crossings for each animal [92]. These data were used to calculate sleep information by extracting immobility bouts of 5 minutes using the *Drosophila* Counting Macro [93,94]. For experiments examining the effects of starvation on sleep, flies were kept on 12:12 LD cycle. Female flies were briefly anesthetized with CO_2_ and placed into plastic tubes containing standard food. All flies were given 24 hours to recover after being anesthetized. Activity was recorded for 24 hours on food, prior to transferring flies into tubes containing 1% agar diluted in dH_2_O (Fisher Scientific) at ZT0. Activity was monitored for an additional 24 hours on agar. For the screen, % change in sleep during starvation was calculated as the sleep duration on agar minus the sleep duration on food, divided by the sleep duration on food for each fly assayed multiplied by a hundred [12,45].

### Immunohistochemistry

The brains of five to seven day old female flies were dissected between ZT4-ZT9 in ice-cold phosphate buffered saline (PBS) and fixed in 4% paraformaldehyde, PBS, 0.5% Triton-X for 30 minutes as previously described [95]. Brains were then rinsed 3X with PBS, 0.5% Triton-X (PBST) for 10 minutes and overnight. In the following day, brains were incubated for 24 hours in primary antibody (1:1000 rabbit anti-Lk; [96] and mouse 1:20 nc82; Iowa Hybridoma Bank) diluted in PBST at 4°C. Brains were rinsed in PBST, 3X for 10 minutes and placed in secondary antibody (1:400 donkey anti-rabbit Alexa 555 and 1:200 donkey anti-mouse Alexa 64; Thermo Scientific), diluted in PBST for 90 minutes at room temperature. Finally, all samples were washed in PBST for a total of 120 minutes and mounted in Vectashield (VectorLabs). Samples were imaged in 2μm sections with a Nikon A1R confocal microscope (Tokyo, Japan) using 20X or 60X oil immersion objective. Images were then processed with NIS Elements 4.40 (Nikon).

### Functional imaging of Lk neurons

Five to seven-day old female flies were collected and placed on vials containing fresh food (Fed) or a wet KimWipe paper (Starved) for 24 hours. All experiments were done between ZT4-ZT7 to account for rhythmic excitability of Lk neurons [25]. For imaging brain explants, previously established methods for calcium imaging were used with modifications [39,69]. Brains of fed or 24hr starved flies were dissected and placed in glass wells (Pyrex) containing artificial hemolymph (140mM NaCL, 2mM KCl, 4.5mM MgCl2, 1.5mM CaC2, and 5mM HEPES-NaOH with Ph=7) and allowed a 5-minute recovery period before being recorded. For 2-Deoxy-D-glucose (2-DG) experiments, fed brains were dissected and placed in 400mM 2-DG (Sigma Aldrich) in artificial haemolymph, 200mM glucose (Sigma Aldrich) in artificial hemolymph, or artificial hemolymph alone for a total of 70 minutes. Every 20 minutes solutions were exchanged. Cover slips were treated with poly-L-lysine (Sigma Aldrich) to ensure that brains were in the same position during imaging and placed onto chamber (RC-21BBDW, Warner Instruments). Fly brains were bathed in artificial hemolymph solution and imaged using a 20X air objective lens on an inverted confocal microscope (Nikon A1R on a Ti-E Inverted Microscope). The pinhole was opened to 244.43μm to allow a thicker optical section to me monitored. UAS-GCaMP-R (GCaMP and mCherry) was expressed in Lk neurons and simultaneously excited with wavelengths of 488nm (FITC) and 561nm (TRITC). Recording were taken for 120 seconds, capturing 1 frame/ 5 seconds with 512 × 512 resolution. For analysis, regions of interest (ROI) were drawn manually, capturing the same area between experimental and control. The mean fluorescence intensity was subtracted from background mean fluorescence intensity for FITC and TRITC per frame. Then, the ratio of GCaMP6.0 to mCherry was calculated and plotted as an average of the total time recorded per brain imaged.

*In vivo* imaging was performed using a previously described protocol with some modifications [97,98]. Briefly, fed or 24 hr starved flies were anesthetized on ice and secured in 200μL pipette tip with head and proboscis accessible. The pipette tip was placed in a small chamber at an angle of 140°, allowing the dorsal and posterior surface of the brain to be imaged. A small hole was cut in the tin foil and fixed to the stage and fly head, leaving a window of cuticle exposed, then sealed using dental glue (Tetric EvoFlow, Ivoclar Vivadent). The proboscis was extended and dental glue was used to secure it in place, ensuring the same position throughout the experiment.

A 21-gauge 1 1/4 needle (PrecisionGlide®, Becton Dickson) was used to cut a window in the fly cuticle. A drop of artificial hemolymph was placed on the cuticle and the connective tissue surrounding the brain was dissected. Flies were allowed to recover from procedure for 30-45 minutes in a humidified box. Mounted flies were placed under a confocal microscope (Nikon A1R on an Upright Microscope) and imaged using a 20X water-dipping objective lens. The pinhole was opened to 244 μm to allow a thicker optical section to be monitored. The settings and data analysis were performed as described above.

### Targeted multi-photon ablation of Lk neurons

Female 3^rd^ instar larvae expressing GFP in Lk neurons were selected and anesthetized in ethyl ether (Fisher Scientific, E134-1) for 2-5 minutes. Larva were placed dorsally on a microscope slide and a cover slip was placed on the larvae. Ringer’s solution was applied onto the larvae below coverslip. Larvae was imaged using a 25X water-dipping objective lens on Multi-photon microscope (Nikon A1R) containing a Chameleon Vision II Ti:Sapphire tunable laser. Excitation laser light of 870 nm was used. Images were acquired at 1 frame per second with a resolution of 512 × 512 pixels. For each neural ablation, a total of 4 frames were acquired. Two frames were captured prior to ablation for a duration of ~2 seconds, followed by ROI stimulation of 2-4 seconds, and 2 frames after ablation. Following ablations, larvae were placed in vials containing food and allowed to grow. Sleep on food and on agar was measured 5-7 days post-eclosion in the *Drosophila* Activity Monitor System. In order to verify which neurons were ablated after behavioral assay, flies were anesthetized on ice and the central nervous system (CNS) was dissected. Fly CNS was fixed in 4% paraformaldehyde, PBS, 0.5% Triton-X for 30 minutes. Following fixation, samples were imaged in 2μm sections with a Nikon A1R confocal microscope (Tokyo, Japan) using 20X oil immersion objective. Ablations that resulted in the formation of supernumerary neurons or deletions of two different subpopulations of Lk neurons were removed from analysis.

### Statistical Analysis

The experimental data are presented as means ± s.e.m. Unless otherwise noted, a one-way or two-way analysis of variance (ANOVA) followed by Tukey’s pot-hoc test was used for comparisons between two or more genotypes and one treatment and two or more genotypes and two treatments. Unpaired t-test was used for comparisons between two genotypes. All statistical analysis were performed using InStat software (GraphPad Software 6.0) with a 95% confidence limit (*p* < 0.05).

---

**S1 Fig.**
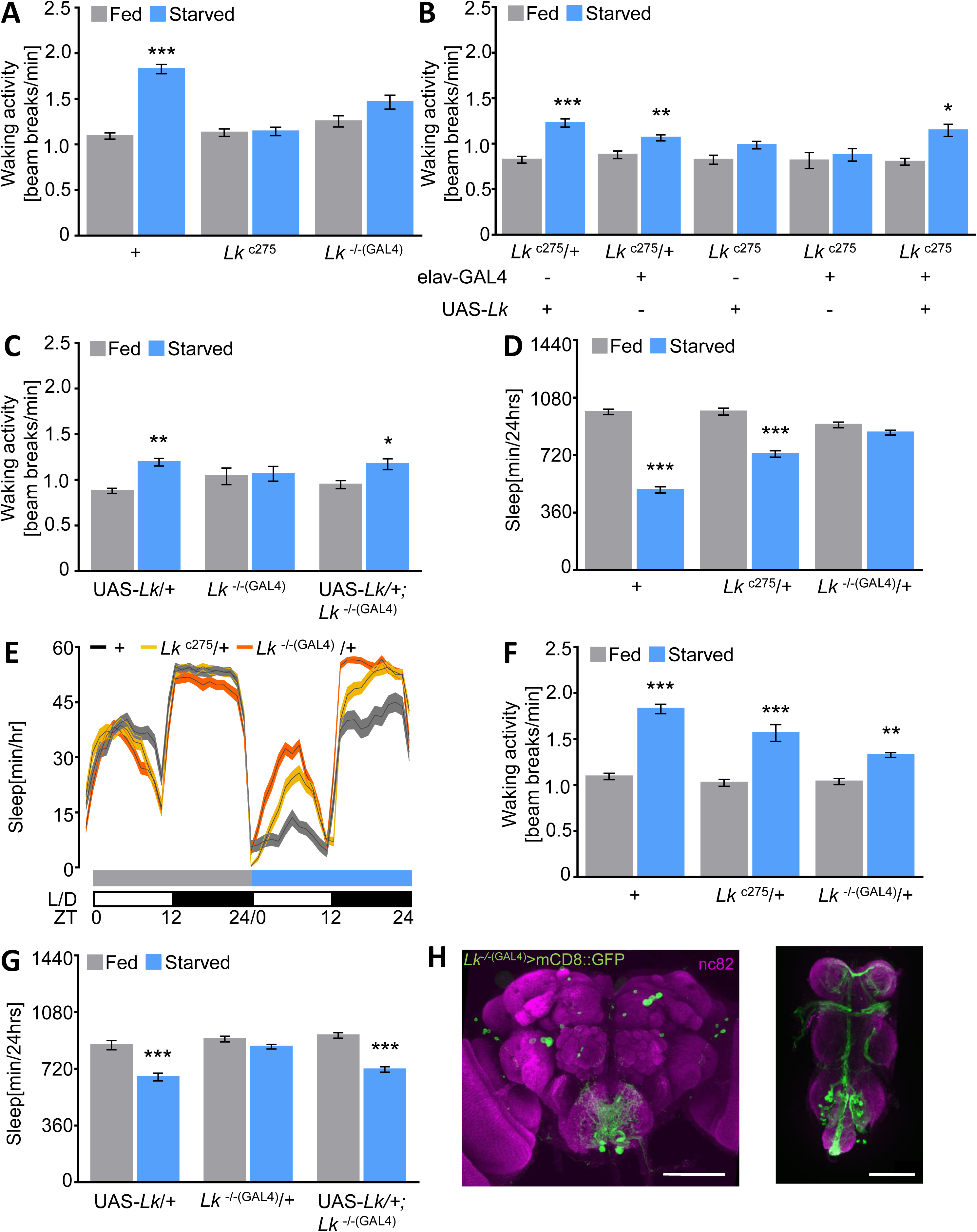
Leucokinin neuropeptide is required for metabolic regulation of sleep. **(A)** Average waking acitivity in fed and starved (blue) flies over 24 hours. Control flies (*w^1118^*) increase waking activity during starvation (n=140, *p*<0.0001) while waking activity does not differ between fed and starved states in *Lk^c275^* (n=58, p=0.99) and *Lk^-/-^*^(GAL4)^ (n≥47, *p*=0.24), Two-way ANOVA (F(2,486)=21.01). **(B)** Pan-neuronal rescue of *Lk^c275^* (elav-GAL4;*Lk^c275^*>UAS-*Lk*;*Lk^c275^*) (n=17, *p*=0.02) restores starvation-induced increase in waking activity compared to *Lk^c275^* mutant controls UAS-*Lk/+*;*Lk^c275^*, (n=23, *p*=0.60) and elav-GAL4/+;*Lk^c275^* (n=20, *p*>0.99). No significant differences were seen during starved state between control flies UAS-*Lk/+*;*Lk^c275^*/+ (n=30, *p*>0.99) or elav-GAL4/*+*;*Lk^c275^*/+ (n=51, *p*>0.99) and rescue flies, Two-way ANOVA (F(4,272)=2.92). **(C)** Increase in waking acitivity following starvation is recued in flies expressing UAS-*Lk* under control of *Lk^-/-^*^(GAL4)^ (n=30, *p*=0.02) compared to *Lk^-/-^*^(GAL4)^ mutants(n=15, *p*>0.99). Flies harboring UAS-Lk alone (UAS-*Lk*/+) increased waking activity following starvation (n=21, *p*=0.02). There were no significant differences during fed state between UAS-*Lk*/+ and rescue (*p*=0.99), or *Lk^-/-^*^(GAL4)^ (*p*=0.69), Two-way ANOVA (F(2,126)=3.09). **(D)** Sleep is significantly reduced in starved *w^1118^* controls (n=140, *p*<0.0001) and flies harboring one copy of *Lk^c275^* (n≥66, *p*<0.0001), while no significant differences are observed in flies harboring one copy of *Lk^-/-^*^(GAL4)^ (n=70, *p*=0.65), Two-way ANOVA (F(2,550)=56.76). **(E)** Sleep profile for hourly sleep averages over a 48 hour experiment on food (day 1, grey) and starved (day 2, blue). Sleep does not differ between any of the groups during day 1. *Lk^-/-^*^(GAL4)^ /+ (orange) have increased sleep on agar during day 2, while wild-type (dark grey) and *Lk^c275^*/+ (gold) suppress sleep. **(F)** *C*ontrols flies (*w^1118^*^)^) (n=140, *p*<0.0001), flies harboring one copy of *Lk^c275^* (n≥66, *p*<0.001) or *Lk^-/-^*^(GAL4)^ alone (n≥70, *p*=0.005) show a significant increase in waking activity during starvation compared to fed state, Two-way ANOVA (F(2,548)=9.19). **(G)** Genetic rescue (UAS-*Lk*/+;*Lk^-/-(^*^GAL4)^/+) (n=86, *p*<0.0001) restores starvation induced suppression compared to flies harboring one copy of *Lk^-/-^*^(GAL4)^ (n=70, *p*=0.40). No significant differences were observed during starved state between heterozygous rescue flies and control flies harboring a copy of UAS-*Lk* alone (n=21, *p*=0.95), Two-way ANOVA (F(2,348)=8.192). **(H)** Whole-brain and ventral nerve cord confocal reconstruction of *Lk^-/-^*^(GAL4)^ >CD8:GFP. GFP-expressing neurons (green) in the brain labeled one pair of neurons in the subeosphogeal zone, four pairs of neurons in the lateral horn, and three pairs of neurons in the medial protorecebrum with variable expression (ALK). In the VNC, Lk is expressed in ~10 pairs of neurons (abdominal Lk neurons; ABLKs). The brain and ventral nerve chord were counterstained with the neuropil marker nc82 (magenta). Scale bar = 100μm. All columns are mean ± SEM; \**p*<0.05; \*\**p*<0.01; \*\*\**p*<0.001.

**S2 Fig.**
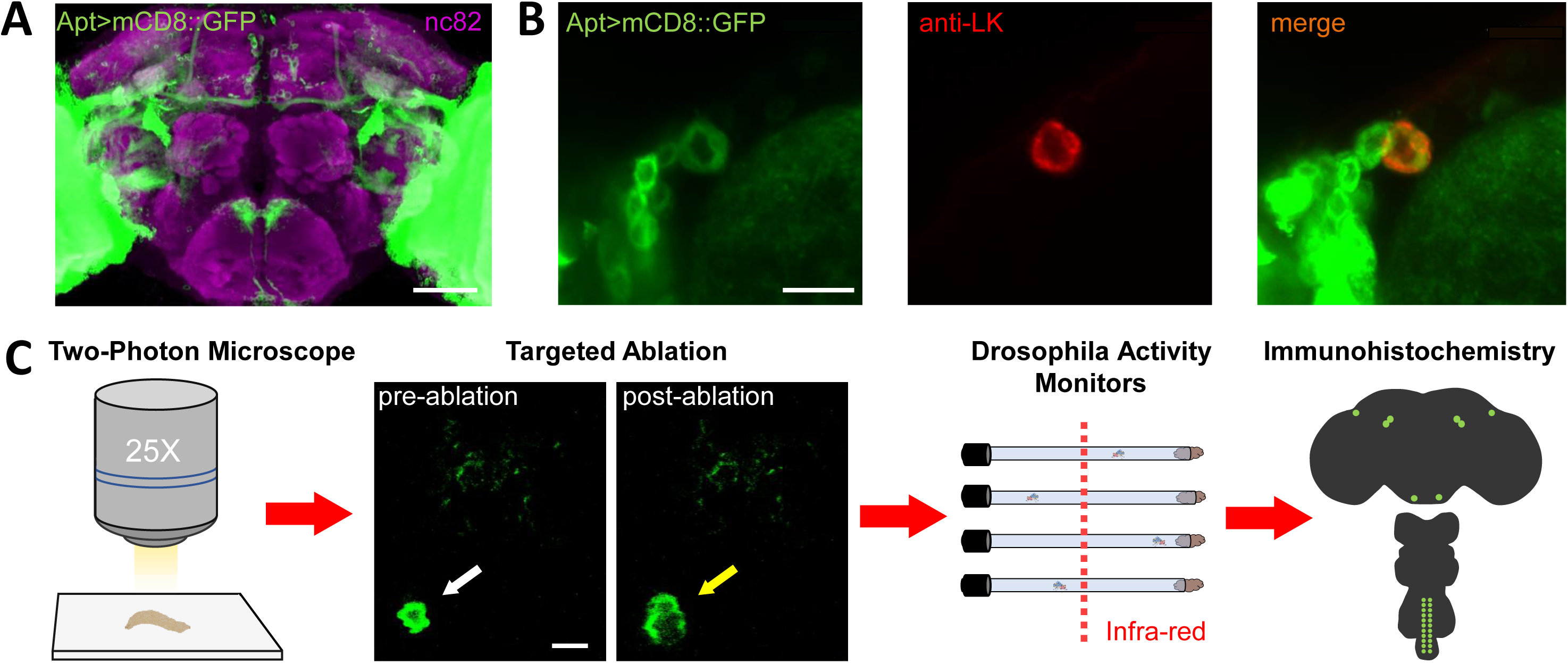
Lateral horn leucokinin neurons are necessary for the metabolic regulation of sleep. **(A)** Expression pattern of Apt-GAL4 driving mCD8::GFP (green). The brain was counterstained with the neuropil marker nc82 (magenta). Scale bar = 50μm. **(B)** Immunostaining for anti-LK (red) in Apt-GAL4>mCD8::GFP (green) reveals LHLK localizes to neurons labelled by Apterous-GAL4 (orange). Depicted is a 14 μm section from the lateral horn region using a 60X oil immersion objective. Scale bar=10μm **(C)** Diagram representative of targeted multi-photon ablation. 3^rd^ instar larvae expressing UAS-mCD8::GFP in Lk neurons are placed drosally onto microscope slide. Two frames are captured prior to ablation (white arrow), followed by ROI stimulation, and 2 additional frames are captured after ablation, where GFP dispersion can be observed (yellow arrow). Following ablation, larvae are placed in vials containing food and allowed to grow. Sleep on food and on agar was measured 5-7 days post-eclosion in the *Drosophila* Activity Monitor System (DAMS). Lastly, immunohistochemistry is performed to verify ablated neurons. Scale bar=10μm.

**S3 Fig.**
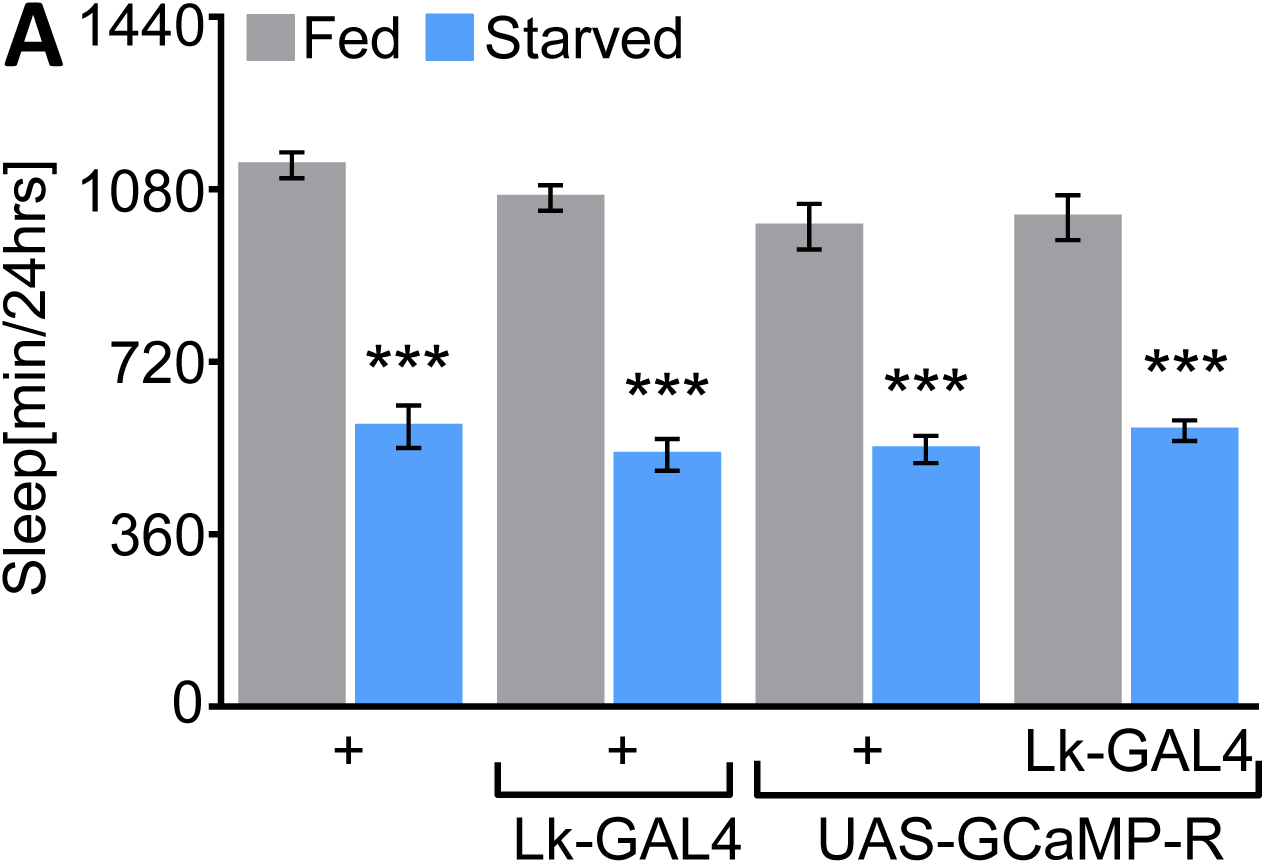
Lateral horn leucokinin neurons have increased activity during starved state. **(A)** Flies expression of UAS-GCaMP-R under Lk-GAL4 sleep significantly more on food (grey) than when starved (blue, n=24, *p*<0.0001) similar to control flies, UAS-GCaMP-R/+ (n=32, *p*<0.0001), Lk-GAL4/+ (n=31, *p*<0.0001), or *w^1118^* flies (n=32, *p*<0.0001). No significant differences were detected on food between Lk-GAL4>UAS-GCaMP-R and *w^1118^* control (*p*=0.45), UAS-GCaMP-R alone (p>0.99) or Lk-GAL4 alone (*p*=0.99), Two-way ANOVA, (F(3,230)=0.97). All columns are mean ± SEM; \*\*\**p*<0.001.

